# P2Y2 purinergic receptor is induced following human cytomegalovirus infection and its activity is required for efficient viral replication

**DOI:** 10.1101/051391

**Authors:** Saisai Chen, Thomas Shenk, Maciej T. Nogalski

## Abstract

Human cytomegalovirus (HCMV) manipulates many aspects of host cell biology to create an intracellular milieu optimally supportive of its replication and spread. The current study reveals a role for purinergic signaling in HCMV infection. The levels of several components of the purinergic signaling system, including the P2Y2 receptor, were altered in HCMV-infected fibroblasts. P2Y2 receptor RNA and protein are strongly induced following infection. Pharmacological inhibition of receptor activity or knockdown of receptor expression markedly reduced the production of infectious HCMV progeny. When P2Y2 activity was inhibited, the accumulation of most viral RNAs tested and viral DNA was reduced. In addition, the level of cytosolic calcium within infected cells was reduced when P2Y2 signaling was blocked. The HCMV-coded UL37x1 protein was previously shown to induce calcium flux from the smooth endoplasmic reticulum to the cytosol, and the present study demonstrates that P2Y2 function is required for this mobilization. We conclude that P2Y2 supports the production of HCMV progeny, possibly at multiple points within the viral replication cycle that interface with signaling pathways induced by the purinergic receptor.

**Importance:** HCMV infection is ubiquitous and can cause life-threatening disease in immunocompromised patients, debilitating birth defects in newborns, and has been increasingly associated with a wide range of chronic conditions. Such broad clinical implications result from the modulation of multiple host cell processes. This study documents that cellular purinergic signaling is usurped in HCMV-infected cells and that the function of this signaling axis is critical for efficient HCMV infection. Therefore, we speculate that blocking P2Y2 receptor activity has the potential to become an attractive novel treatment option for HCMV infection.

## Introduction

Human cytomegalovirus (HCMV) is a beta herpesvirus that infects a large percentage of the adult population worldwide. Infection in immunocompetent people is typically asymptomatic. In contrast, HCMV is a leading opportunistic pathogen in immunosuppressed individuals [1–4] and is a major infectious cause of birth defects [5, 6].

HCMV causes broad cellular effects that likely contribute to the diverse pathologies associated with infection. One mechanism utilized by the virus to change the biology of infected cells is via the regulation of expression levels and activities of cell surface proteins [7, 8]. Here, we describe the role of purinergic receptors during HCMV infection. Purinergic receptors are ubiquitous cell surface receptors that are activated by extracellular adenosine di-(ADP) and tri-(ATP) phosphates (P2 receptors) or adenosine (P1 receptors). P2 purinergic receptors are further divided into ionotropic P2X and metabotropic P2Y families. P2X receptors are ATP-gated ion channels and P2Y receptors are G protein-coupled receptors that are activated by adenine and uridine nucleotides or nucleotide sugars. Seven subtypes of P2X receptors (P2X1-7) and eight of P2Y receptors (P2Y1, P2Y2, P2Y4, P2Y6, P2Y11, P2Y12, P2Y13, and P2Y14) have been identified [9, 10].

Signaling via the P2Y receptors triggers the activation of a heterotrimeric G protein, which leads to the activation of effector protein phospholipase C (PLC), and generation of diacylglycerol (DAG) and inositol 1,4,5-triphophosphate (IP_3_)[11]. DAG stimulates protein kinase C (PKC)[12]. The activation of PKC has been shown to increase the expression of early growth response protein 1 (Egr-1) [13], a transcription factor critical for DNA synthesis, proliferation, and migration of fibroblasts and other cells [14]. IP_3_ mobilizes cytosolic Ca^2+^ from the smooth endoplasmic reticulum (SER). It has been observed that in lung fibroblasts, P2Y2 is the only purinergic receptor subtype that, when activated, causes the mobilization of intracellular Ca^2+^ [15]. P2Y2-mediated intracellular Ca^2+^ increases have been implicated in promoting the proliferation and migration of hepatocellular carcinoma cells in mice [16]. Moreover, P2Y2 signaling was found to stimulate HIV-1 viral fusion through the activation of proline-rich tyrosine kinase 2 (Pyk2) [17].

P2X receptors differ notably from their P2Y counterparts in ligand selectivity. While P2Y receptors recognize a wide range of agonists, P2X receptors are activated only by ATP. Moreover, P2X receptors are assembled as trimeric proteins. Specifically, P2X5 mainly functions as heterotrimers involving P2X1, P2X2, or P2X4 [18]. The expression of P2X5 receptors is normally restricted to the trigeminal mesencephalic nucleus of the brainstem, sensory neurons, cervical spinal cord, and some blood vessels [19]. P2X5 receptor expression in atypical locations has been linked to cancer [20–22]. Activation of the P2X5 receptor causes an influx of cations (Na^+^, K^+^, and Ca^2+^) across the plasma membrane [23].

Knowing that purinergic receptors regulate cellular calcium levels and that an increase in calcium levels has been observed following HCMV infection [24], it is intriguing to speculate that members of the family of purinergic receptors could be important factors during HCMV infection. It has been reported previously that a few members of the family of purinergic receptors exhibited higher expression in HCMV-infected cells, however their molecular role during infection has not been found [25, 26]. The regulation of Ca^2+^ release is critical for viral DNA synthesis and the production of infectious progeny [27]. It has been found that the HCMV immediate early protein pUL37x1 induces the mobilization of Ca^2+^ from SER to the cytosol [28]. Altering the release of Ca^2+^ from SER affects the activity of Ca^2+^-dependent ER chaperones, resulting in the accumulation of unfolded proteins and contributing to the unfolded protein response [29]. In addition, the Ca^2+^-dependent protein kinase, PKCα, is activated following infection, leading to the production of large (1-5 μm diameter) cytoplasmic vesicles at late times during infection [30]. The presence of these vesicles correlates with the efficient accumulation of enveloped virions. The release of Ca^2+^ from SER during infection is also expected to influence Ca^2+^-dependent processes that occur in the mitochondria. For instance, Ca^2+^ uptake can stimulate aerobic metabolism and enhance ATP production [31, 32]. Ca^2+^ can also induce the activity of Ca^2+^/calmodulin-dependent protein kinase kinase (CaMKK), which activates 5’ AMP-activated protein kinase (AMPK). AMPK activity has been shown to support HCMV-induced changes to the infected cell metabolome, as well as to be necessary for HCMV DNA synthesis and the expression of viral late genes [33, 34]. Interestingly, blocking viral DNA synthesis by inhibiting viral DNA polymerase, pUL54, can serve as an effective therapeutic measure for HCMV infection [35]. Additionally, cellular phosphoinositide 3-kinase (PI3-K) and p38 kinase activities were found to be required for viral DNA replication and the production of infectious progeny [36, 37]. As both PI3-K and p38 are downstream factors of the purinergic receptor signaling axis [38, 39], it is conceivable that purinergic receptors can have an effect on viral DNA synthesis. Finally, since Ca^2+^ plays a central role in mediating apoptosis, regulating the concentration of intracellular Ca^2+^ may affect the infected cell’s sensitivity to apoptotic stimuli [40, 41].

Although the viral pUL37x1 protein has already been implicated in mediating Ca^2+^ release during HCMV infection, there may also be other pathways that affect intracellular Ca^2+^ levels. Therefore, we speculated that the purinergic receptor-mediated signaling may work alongside pUL37x1 in regulating Ca^2+^ release into the cytosol, influencing the efficiency of HCMV infection. In fact, one study using mouse fibroblasts showed that extracellular ATP and UTP could stimulate the mobilization of Ca^2+^ from intracellular stores by an IP_3_-mediated pathway downstream of P2Y2 signaling that is independent of PKC activation [42]. To test this notion, we used pharmacological agents and siRNA technology to study the consequences of inhibiting purinergic receptors during HCMV infection. We provide evidence that P2Y2 inhibition interferes with the expression of viral genes and the release of infectious progeny, whereas P2X5 inhibition enhances HCMV yield. P2Y2-mediated signaling affects HCMV infection at the stage of viral DNA synthesis and significantly contributes to the regulation of intracellular Ca^2+^ homeostasis.

## Methods and Materials

### Cells, viruses and drugs

Human foreskin fibroblasts (HFF) and human lung fibroblasts (MRC5) were cultured in Dulbecco’s Modified Eagle Medium (DMEM; Sigma-Aldrich) supplemented with 10% fetal bovine serum (FBS). Penicillin and streptomycin were added to the media for all experiments except those involving siRNA transfection.

HFF cells stably expressing the viral protein, pUL37x1, were made using the pLVX-EF1α promotor lentivirus packaging plasmid (Clontech). To insert AD169 UL37x1 between EcoRI-BamHI sites of the pLVX-EF1 plasmid, the sequence was amplified by PCR from BADwt, a BAC containing the AD169 strain [43]. Lentivirus particles were generated using sequence-confirmed plasmids containing UL37x1 or GFP as a control. Following lentiviral treatment, cells expressing pUL37x1 or GFP were selected using 2 μg/ml puromycin, and protein expression was evaluated by Western blot using mouse monoclonal antibody 4B6-B [28].

Previously described HCMV strains TB40/E, TB40-GFP, AD*wt* and ADsubUL37x1 HCMV strains were used in these studies [43, 44]. For some experiments, TB40/E virus was inactivated by irradiation (50 J/m^2^) in a Stratalinker (Stratagene) [45]. A pUL37x1-deficient derivative of AD*wt*, AD*sub*UL37x1 [43], lacks the AD169 genomic sequence from 169,144 to 169,631 and includes kanamycin resistance and *LacZ* markers [46]. All viral stocks were purified by centrifugation through a 20% D-sorbitol cushion containing 50mM Tris·HCl, 1mM MgCl_2_, pH 7.2, and resuspended in DMEM. Infections were performed by treating cells with viral inoculum for 2 h, followed by removal of inoculum and washing with phosphate-buffered saline (PBS) before applying with fresh media.

Virus titers were determined based on viral IE1 expression on a reporter plate containing fibroblasts [34, 47]. At 24 hpi, cells were assayed by indirect immunofluorescence using a primary antibody against the viral IE1 protein [48] and secondary goat anti-mouse antibody conjugated to Alexa Fluor 488 (Molecular Probes). Nuclei were stained using Hoescht dye (Thermo Fisher Scientific). IE1-positive nuclei were quantified using the Operetta High-Content Imaging System (PerkinElmer). In some cases, titers were determined by TCID_50_ analysis, and calculations were performed according to the Reed and Muench formula [49]. All virus stocks were titered by the TCID_50_ method.

Kaempferol (Sigma-Aldrich) was dissolved in DMSO and stored at 4°C until use. PPADS tetrasodium salt (Torcis) was dissolved in dimethyl water and stored at −20°C until use. Cytotoxicity assays for drugs were performed at 96 h post treatment using the CellTiter 96^®^ AQueous One Solution Cell Proliferation Assay (Promega, Madison, WI), according to the manufacturer’s protocol. Cell viability was measured based on absorbance at 490 nm using a SpectraMax Plus spectrometer (Molecular Devices).

### siRNA knockdown experiments

siRNAs targeting P2Y2 (siP2Y2), P2X5 (siP2X5), ENPP4 (siENPP4) or ENTPD2 (siENTPD2) were designed and purchased from Life Technologies. HFFs were grown to ~80% confluence in DMEM supplemented with 10% FBS and then transfected with siRNA using Lipofectamine^®^ RNAiMAX Reagent (Life Technologies). Cells transfected with non-specific, scrambled siRNA (siSc) (Life Technologies) served as a negative control. Following a 24 h-incubation, cells were washed with PBS and either infected with TB40/E-GFP virus or mock infected.

### Assay for viral entry

For drug assays, kaempferol, PPADS, or solvent was applied to 95-100% confluent HFFs for 1 h, and for siRNA analyses, cells were transfected with siP2Y2, siP2X5 or siSc for 24 h. Cells were then washed with PBS and infected at a multiplicity of 1 TCID_50_/cell or mock infected for 1 h at 4°C. Virus that had not penetrated cells was removed with a low-pH citrate buffer (40 mM citric acid, 10 mM KCl, 135 mM NaCl, pH 3.0) [50]. Cells were then incubated at 37°C in DMEM containing 10% FBS. Cells were either fixed in cold MeOH and immunostained for viral IE1 protein at 24 hpi, as described above, or viral DNA was isolated 1 h after the reaction temperature was raised to 37°C, using the QIAamp DNA Mini kit (Qiagen) and quantified by quantitative polymerase chain reaction (qPCR) performed in duplicate using SYBR Green master mix (Applied Biosystems) using the QuantStudio^™^ 6 Flex Real-Time PCR System (Applied Biosystems). Primers for the viral UL123 locus were used (Table 1). Fold change was calculated using the ΔΔCt method and GAPDH was used as an internal control [51].

**TABLE 1:**
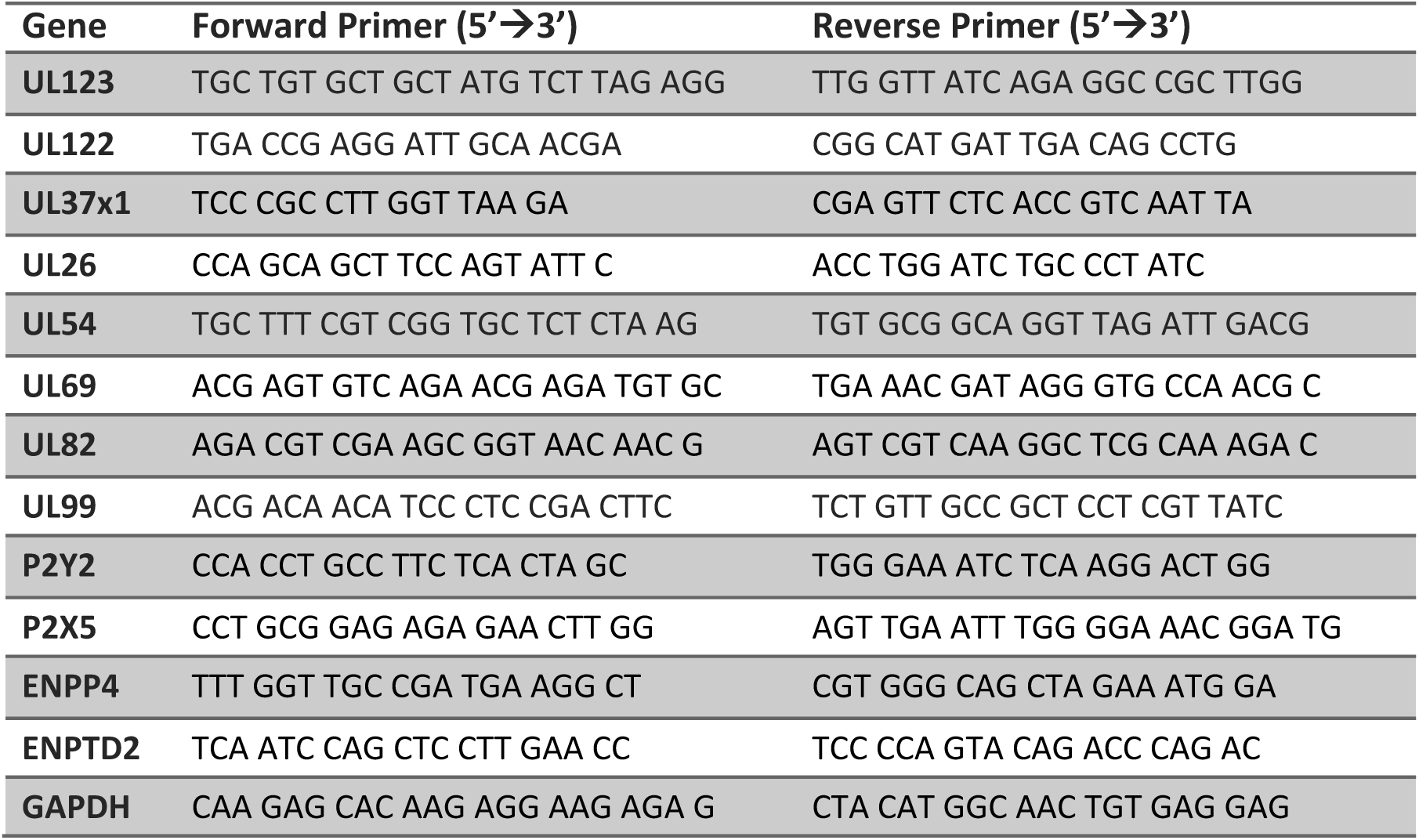
Primers used in qPCR assays.

### Nucleic acid and protein analyses

For RNA-Seq analysis of protein-coding RNAs, MRC-5 human lung fibroblasts were infected with the AD169 strain of HCMV. RNA from mock-and HCMV-infected cells was collected at 48 hpi and isolated using the miRNeasy kit (Qiagen.). RNA quality was analyzed using the Bioanalyzer 2100 (Agilent Technologies). TruSeq Stranded Total RNA Library Prep Kit with Ribo-Zero (Illumina) was used to generate cDNA libraries as per the manufacturer’s instructions. Briefly, cytosolic ribosomal RNA (rRNA) was depleted from total RNA using biotinylated probes that selectively bind rRNA species. The resulting rRNA-depleted RNA underwent fragmentation, reverse transcription, end repair, 3’-end adenylation, adaptor ligation and subsequent PCR amplification and SPRI bead purification (Beckman Coulter). The unique barcode sequences were incorporated in the adaptors for multiplexed high-throughput sequencing. The final product was assessed for its size distribution and concentration using BioAnalyzer High Sensitivity DNA Kit (Agilent) and Kapa Library Quantification Kit (Kapa Biosystems). The libraries were pooled, denatured and loaded onto a HiSeq Rapid Paired-Read flow cell (Illumina), which was subjected to 2X100 cycles of sequencing by an Illumina HiSeq 2500 (Illumina). Illumina CASAVA pipeline Version 1.8 was used to extract de-multiplexed sequencing reads. Sequencing reads were normalized using the trimmed mean of M-values method (TMM), mapped to the human reference genome and transcripts annotated to which reads have been mapped. Mapped reads were counted and the differential gene expression between triplicate samples from mock-and HCMV-infected cells was computed based on counts per million (CPM). Ingenuity Pathway Analysis cloud software (Qiagen) was used to group the genes identified as significantly up-or down-regulated into gene ontologies and to determine which cellular networks were the most significantly regulated in HCMV-infected cells.

For quantitative reverse transcription PCR (qRT-PCR) analysis of RNA, RNA was extracted from samples collected in QIAzol lysis reagent using the miRNeasy kit (Qiagen). Complimentary DNA (cDNA) was made from 1 μg of total RNA with oligo dT and MultiScribe reverse transcriptase (Applied Biosystems), according to the manufacturer’s protocol. Reverse transcription reactions were run at 25°C for 10 min, 48°C for 30 min, and 90°C for 5 min. Primers used in the study are listed in Table 1. qPCR was performed in duplicates on equal volumes of cDNA using SYBR Green master mix (Applied Biosystems) on the QuantStudio^™^ 6 Flex Real-Time PCR System (Applied Biosystems). Transcript levels were analyzed using the ΔΔCt method and GAPDH was used as an internal control [51]. Error ranges are reported as standard error of the mean (SEM).

For quantification of viral DNA, total DNA was isolated from HCMV-infected cells or media at 96 hpi, and viral DNA was quantified by qPCR using primers specific for UL123 gene and a standard curve created by serially diluting BAD*wt* DNA. To calculate the particle-to-infectious unit ratio, media collected at 96 hpi was titered and viral DNA was isolated from DNase I-treated (10U/mL; 30 min. at 37C) virions to calculate viral DNA copy number/mL as a measure of virus particles.

For protein analysis, fibroblasts were harvested using lysis buffer (50 mM Tris-HCl, pH 7.5, 5 mM EDTA, 100 mM NaCl, 1% Triton X-100, 0.1% SDS, and 10% glycerol). Samples were mixed with 6x SDS sample buffer (325 mM Tris pH 6.8, 6% SDS, 48% glycerol, 0.03% bromophenol blue containing 9% 2-mercaptoethanol). Equal protein amounts of the different samples were separated by electrophoresis (SDS-PAGE) and transferred to ImmunoBlot polyvinylidene difluoride (PVDF) membranes (BioRad Laboratories). Western blot analyses were performed using primary antibodies recognizing P2Y2 (H-70; Santa Cruz Biotechnology), P2X5 (Sigma-Aldrich), β-actin-HRP (Abcam), or pUL37x1-specific mouse monoclonal antibody 4B6-B [28]. Goat anti-mouse and donkey anti-rabbit (GE Healthcare Biosciences) conjugated with horseradish peroxidase (HRP) were used as secondary antibodies. Western blots were developed using WesternSure ECL Detection Reagents (Licor).

### Intracellular calcium assay

HFFs were cultured to 95% confluency, drug-treated and infected at a multiplicity of ~1 with *ADwt*, AD*sub*UL37x1 viruses or mock infected. At 20 hpi, cells were washed and incubated in media containing 2 μM Fluo-4 AM dye (Life Technologies) for 30 min. at 37°C. After washing with PBS, images were captured using the Operetta High-Content Imaging System (PerkinElmer) and the fluorescent signal was measured based on 10 cells per experimental arm using ImageJ software [52]. Background fluorescence was subtracted and fold change was determined relative to control samples.

## Results

### HCMV infection elevates the steady-state levels of several purinergic receptors

To investigate the differential expression of purinergic receptors during infection, we performed whole RNA sequencing of MRC-5 fibroblasts infected with the AD169 strain of HCMV. RNA from mock and infected cells was analyzed in three independent experiments where samples were collected at 48 hpi, allowing us to analyze RNAs during the late phase of the viral replication cycle. Several components involved in purinergic receptor signaling are expressed at significantly higher levels in the infected cells, including P2Y2, P2X5, ENPP4 and ENTPD2, and several others were reduced (Table 2).

**TABLE 2:** Differential expression of members of the purinergic receptor network in HCMV-infected MRC-5 fibroblasts at 48 hpi compared to mock-infected controls. Log_2_ fold change (FC) was calculated from sequence reads determined from triplicate biological samples.

| Gene | Description | Log<sub>2</sub> (FC) |
| --- | --- | --- |
| P2Y2 | Purinergic receptor, P2Y subtype 2 | 7.49 |
| ENPP4 | Ectonucleotide pyrophosphatase/phosphodiesterase 4 | 4.20 |
| P2X5 | Purinergic receptor, P2X subtype 5 | 3.21 |
| ENTPD2 | Ectonucleoside triphosphate diphosphohydrolase 2 | 3.18 |
| P2X1 | Purinergic receptor, P2X subtype 1 | 2.78 |
| ENTPD8 | Ectonucleoside triphosphate diphosphohydrolase 8 | 2.65 |
| PANX2 | Pannexin channel 2 | 2.28 |
| ENPP5 | Ectonucleotide pyrophosphatase/phosphodiesterase 5 | 1.41 |
| ENPP6 | Ectonucleotide pyrophosphatase/phosphodiesterase 6 | -3.66 |
| ENPP7P10 | Ectonucleotide pyrophosphatase/phosphodiesterase 7<br>pseudogene 10 | -3.58 |
| P2X6 | Purinergic receptor, P2X subtype 6 | -2.76 |
| ENPP7P4 | Ectonucleotide pyrophosphatase/phosphodiesterase 7<br>pseudogene 4 | -2.17 |
| P2X7 | Purinergic receptor, P2X subtype 7 | -1.97 |
| NT5E | Ecto-5' nucleotidase (CD73) | -1.68 |
| ENPP7P12 | Ectonucleotide pyrophosphatase/phosphodiesterase 7<br>pseudogene 12 | -1.55 |

To confirm the results obtained from RNA-Seq and extend the findings to a different fibroblast cell population and virus strain, we used qRT-PCR to investigate the expression levels of the RNAs identified in our RNA-seq analysis. HFFs were infected with the TB40/E-GFP strain of HCMV at a multiplicity of 3 TCID_50_/cell. At 48 hpi, HCMV-infected cells expressed 76.41 ± 20.10 times more P2Y2, 2.74 ± 0.54 times more P2X5, 9.98 ± 3.22 times more ENPP4 and 45.55 ± 16.54 times more ENTPD2 RNA than mock-infected controls (Fig. 1).

**FIG 1.**
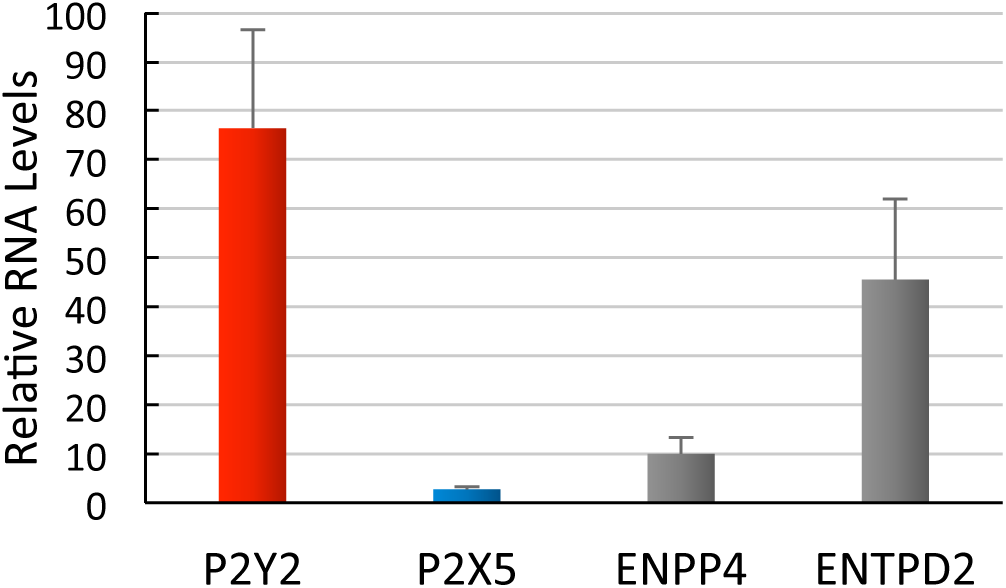
HCMV infection increases the transcript levels of P2Y2, P2X5, ENPP4, and ENTPD2 at 48 hpi. HFFs were infected with TB40/E-GFP vims (MOI=3) or mock infected for 2 h. Samples were collected at 48 hpi, transcripts were quantified by qRT-PCR and GAPDH was used as an internal control. Data are represented as fold change compared to mock-infected cells. Results were averaged between two biological replicates and error bars represent standard error of the mean (SEM).

We also examined the kinetics of P2Y2 and P2X5 receptor expression throughout the viral replication cycle. HFFs were infected at a multiplicity of 3 TCID_50_/cell, and samples were collected after various time intervals. P2Y2 and P2X5 RNAs were assayed by qRT-PCR and receptor proteins were assayed by Western blot. P2Y2 RNA was elevated at 24 hpi and increased as the infection progressed (Fig. 2A). P2Y2 protein became highly abundant at 48 hpi and its levels stayed elevated through the last time point assayed at 120 hpi (Fig. 2C). These results also qualitatively match the mass spectroscopy-based analysis of cell surface proteins, which determined that P2Y2 protein is 3-, 4-and 5-fold more abundant at 24, 48, and 72 hpi, respectively, compared to uninfected cells [8]. In the case of P2X5, its transcript levels increased up to 48 hpi and then decreased later in infection (Fig. 2B). This corresponded with increasing levels of P2X5 protein between 24 and 72 hpi, which then markedly decreased later in infection (Fig. 2C).

**FIG 2.**
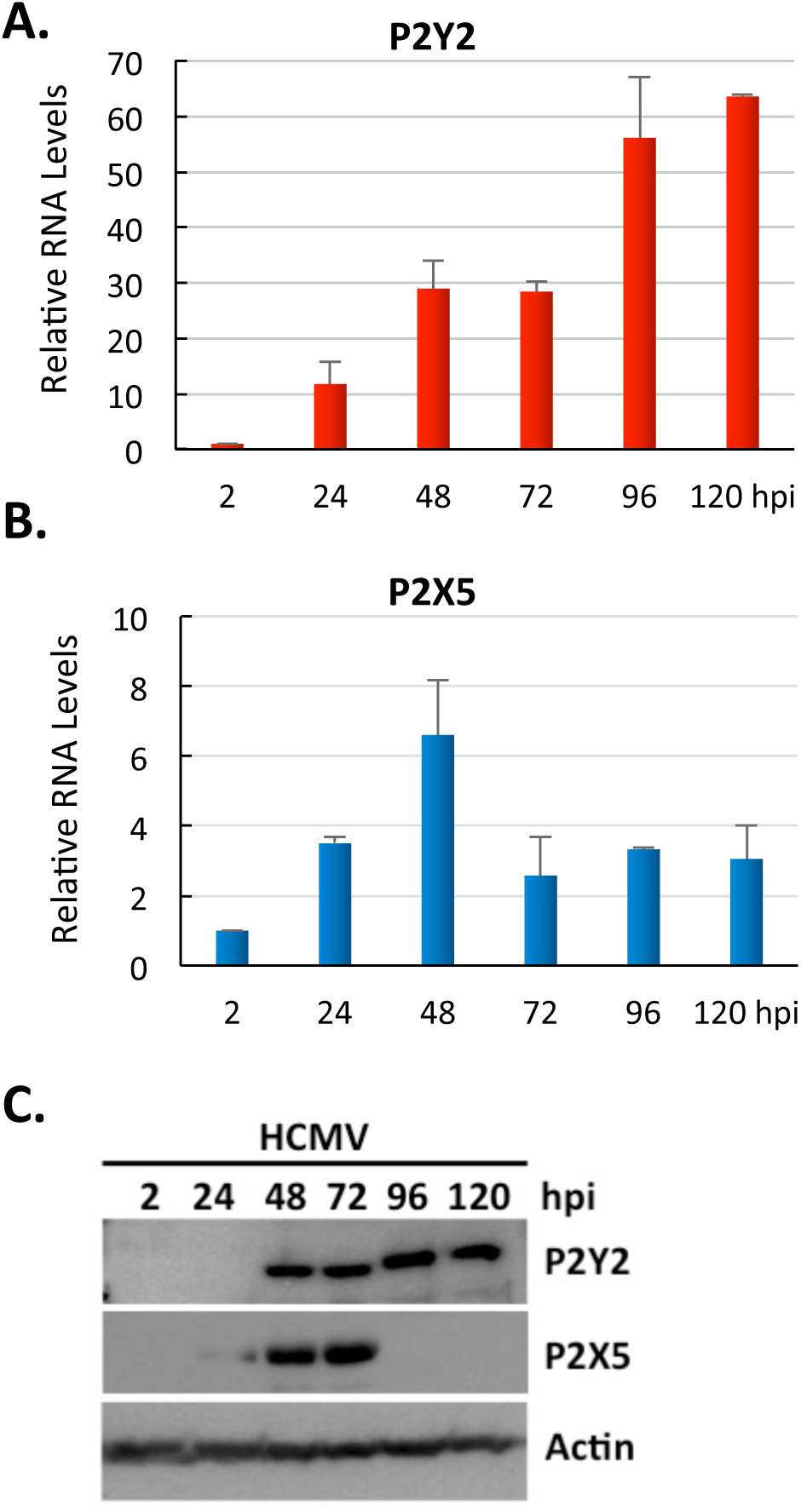
Kinetics of P2Y2 and P2X5 expression in HCMV-infected HFFs. **(A, B) P2Y2 and P2X5 RNA levels are modulated by infection**. HFFs were infected with TB40/E-GFP (MOI=3) or mock infected. RNA and protein samples were collected at various times after infection, transcripts were quantified by qRT-PCR and GAPDH was used as an internal control. Results are shown as fold change compared to mock-infected cells. Data were averaged for two biological replicates and error bars represent SEM **(C) P2Y2 and P2X5 proteins are modulated by infection**. Proteins were subjected to Western blot analyses using antibodies specific for P2Y2 and P2X5. Actin was monitored as a loading control.

Taken together, data from both RNA-Seq and qRT-PCR analyses confirm that HCMV infection in fibroblasts causes an increase in the expression of the cellular purinergic receptors P2Y2 and P2X5, as well as the cell surface ectonucleotidases ENPP4 and ENTPD2.

### Viral gene expression is necessary for the normal modulation of P2Y2 and P2X5 expression

Since the overexpression of these receptors was seen during the early and late phases of infection, we wished to determine whether the binding of the virion to the fibroblast cell surface was sufficient to induce their upregulation. To do so, we used UV-irradiated TB40/E virus that can bind to and enter cells but cannot express its genes. HFFs were infected with the untreated virus or UV-irradiated virus at a multiplicity of 3 TCID_50_/cell, samples were collected at 24 and 48 hpi and receptor transcript levels were quantified by qRT-PCR. Compared to cells infected with untreated TB40/E virus, those infected with the UV-irradiated virus exhibited 90% lower expression of P2Y2 RNA (Fig. 3A) but 40% higher expression of P2X5 (Fig. 3B). This finding suggests that viral binding and entry is not sufficient to produce the increased levels of P2Y2 transcript normally seen during infection. However, viral gene expression appears not to be required for the upregulation of P2X5 expression.

**FIG 3.**
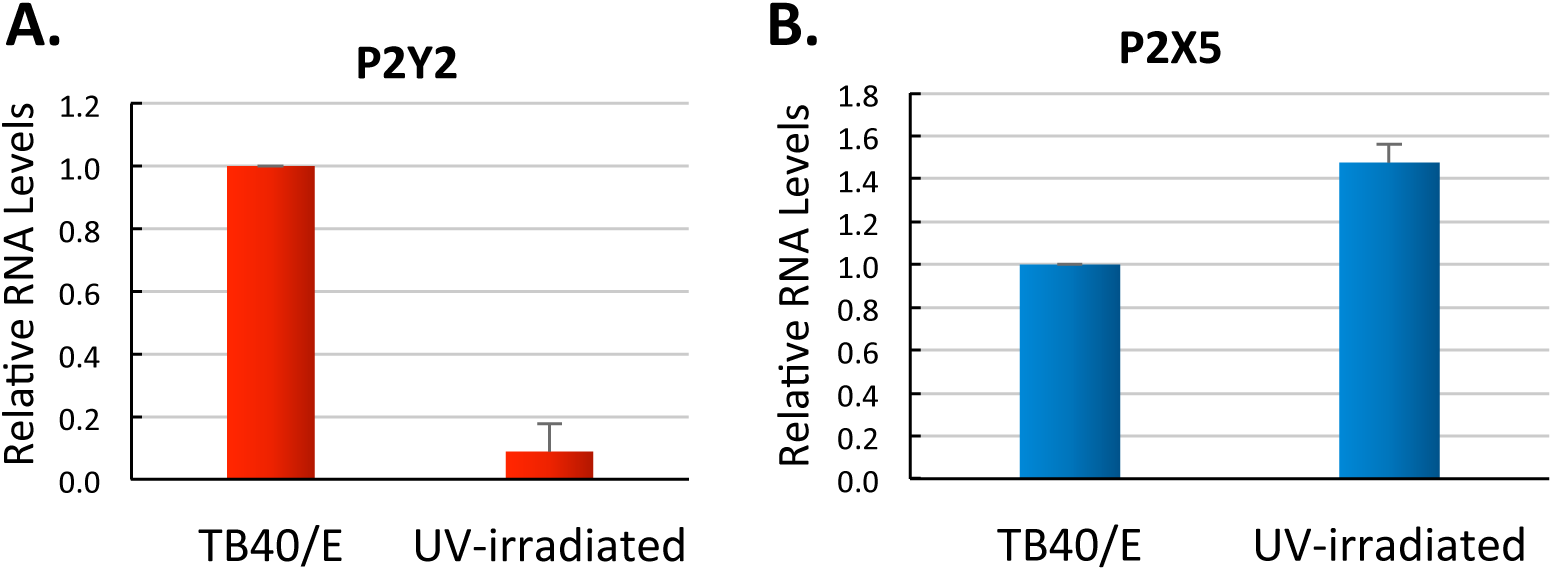
UV-irradiated HCMV failed to increase P2Y2 but did increase P2X5 RNA levels. Untreated or UV-irradiated TB40/E virus was applied to HFFs (MOI=3). Samples were collected at 48 hpi (P2Y2) or 24 hpi (P2X5), and transcripts were quantified by qRT-PCR. GAPDH was assayed as an internal control. Results are shown as fold change compared to untreated control virus condition. Data were averaged between two biological replicates and error bars represent SEM.

### P2Y2 and P2X5 receptors have opposite effects on HCMV yield

To determine whether the upregulated expression of these purinergic receptors and ectonucleotidases is related to HCMV pathogenesis, we assessed the effects of inhibiting their expression or activity on the production of infectious progeny.

First, we employed siRNAs to knock down their expression. Specific siRNAs designed to target the P2Y2 (siP2Y2), P2X5 (siP2X5), ENPP4 (siENPP4) and ENTPD2 (siENTPD2) transcripts were used to inhibit the expression of these four genes. We verified that the siRNAs reduced expression of their targeted transcripts by using qRT-PCR (Fig. 4A). HFFs were transfected with the selected siRNAs for 24 h before HCMV infection at a multiplicity of 3 TCID_50_/ml. At 120 hpi, the media was collected and viral titer assayed. Knockdown of the P2Y2 (~15% of its normal level) or P2X5 receptor (~10% of its normal level) reduced the viral yield by ~95% or increased the yield by almost 400%, respectively (Fig. 4B), compared to viral titer of the media collected from scrambled siRNA (siSc)-treated cells. In contrast, siRNA-mediated knockdown of ENPP4 or ENTPD2 had minimal effects on the amount of infectious progeny released by the infected cells (Fig. 4B).

**FIG 4.**
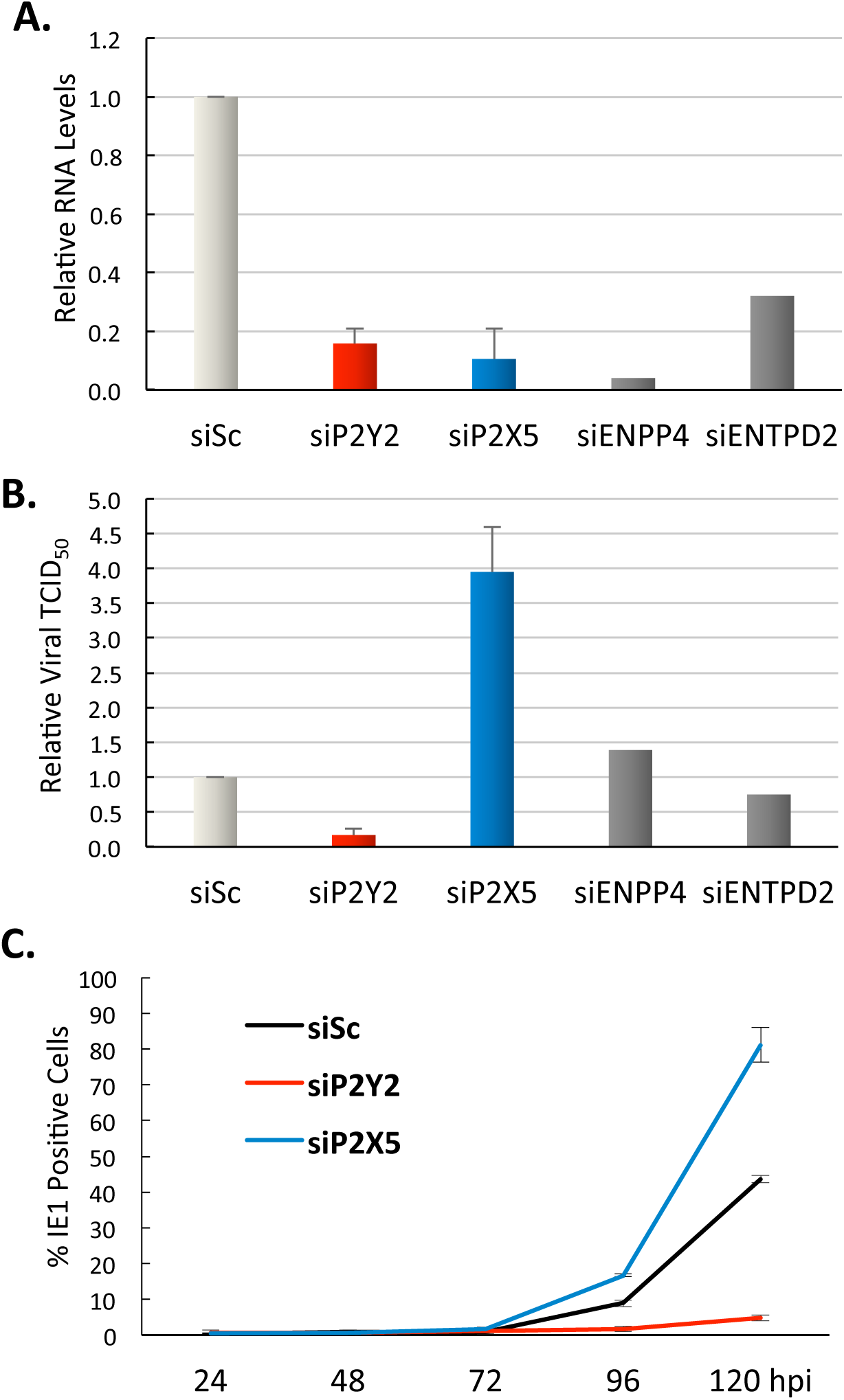
siRNA-mediated knockdown of P2Y2 or P2X5 affects HCMV yield. HFFs were transfected with siP2Y2, siP2X5, siENPP4, siENTPD2, or scrambled siRNA (siSc) as a control. After 24 h, cells were infected with TB40/E-GFP virus (MOI=3). **(A) siRNA validation**. Samples were collected at 48 hpi and qRT-PCR was used to measure levels of P2Y2, P2X5, ENPP4 and ENTPD2 transcripts. GAPDH was used as an internal control. The data are represented as fold change compared to siSc control. The mean fold change for three biological replicates are shown and error bars represent SEM. **(B) Effect of siRNA-mediated knockdowns on virus yield at 120 hpi**. TCID_50_/ml values were determined and fold change was calculated relative to siSc control. Results show the average fold change from two biological replicates and error bars represent SEM. **(C) Virus growth kinetics following siRNA-mediated knockdown**. Samples were collected after various time intervals. Viral titers were determined by applying the infectious media to a reporter plate of HFFs and immunostaining for IE1 protein 24 h later. The data are presented as percent of IE 1-positive cells averaged from two biological replicates and error bars represent SEM.

Seeing the significant effect of P2Y2 and P2X5 knockdown on HCMV yield, we focused further study on the roles of these two receptors. To investigate the kinetics of viral release during infection, HFFs were transfected with selected siRNAs for 24 h before HCMV infection at a multiplicity of 3 TCID_50_/cell. Differences in the amount of virus released from cells transfected with siP2Y2 (reduced virus yield) or siP2X5 (increased virus yield) relative to control cells transfected with siSc were first observed at 72 hpi and gradually increased up to 120 hpi (Fig. 4C).

We next employed pharmacological perturbations to confirm the roles of the receptors during HCMV infection. Kaempferol is a selective P2Y2 receptor antagonist [17, 53]. PPADS has ~10-fold higher affinity for blocking P2X5 than other P2X family members [23, 54]. We used kaempferol and PPADS at 50 μM in our studies, because the drugs are commonly used to treat fibroblasts at concentrations of 20-100 μM [55–58] and we detected little toxicity when uninfected fibroblasts were treated at doses ranging from 0-400 μM (Fig. 5A, B). To test the effects of inhibiting P2Y2 and P2X5 receptor activity on the release of viral progeny, HFFs were pretreated with drug for 1 h prior to HCMV infection at a multiplicity of 3 TCID_50_/cell. After allowing 2 h for viral entry into cells, cells were washed and supplemented with media containing either kaempferol or PPADS. Media with drug was replaced every 24 h until media samples were collected at 96 hpi and assayed for infectious virus. Kaempferol decreased the viral yield by more than 99%, whereas PPADS nearly tripled the yield (Fig. 5C). Therefore, we conclude that elevated P2Y2 receptor activity normally facilitates infection whereas the P2X5 receptor plays an antiviral role.

**FIG 5.**
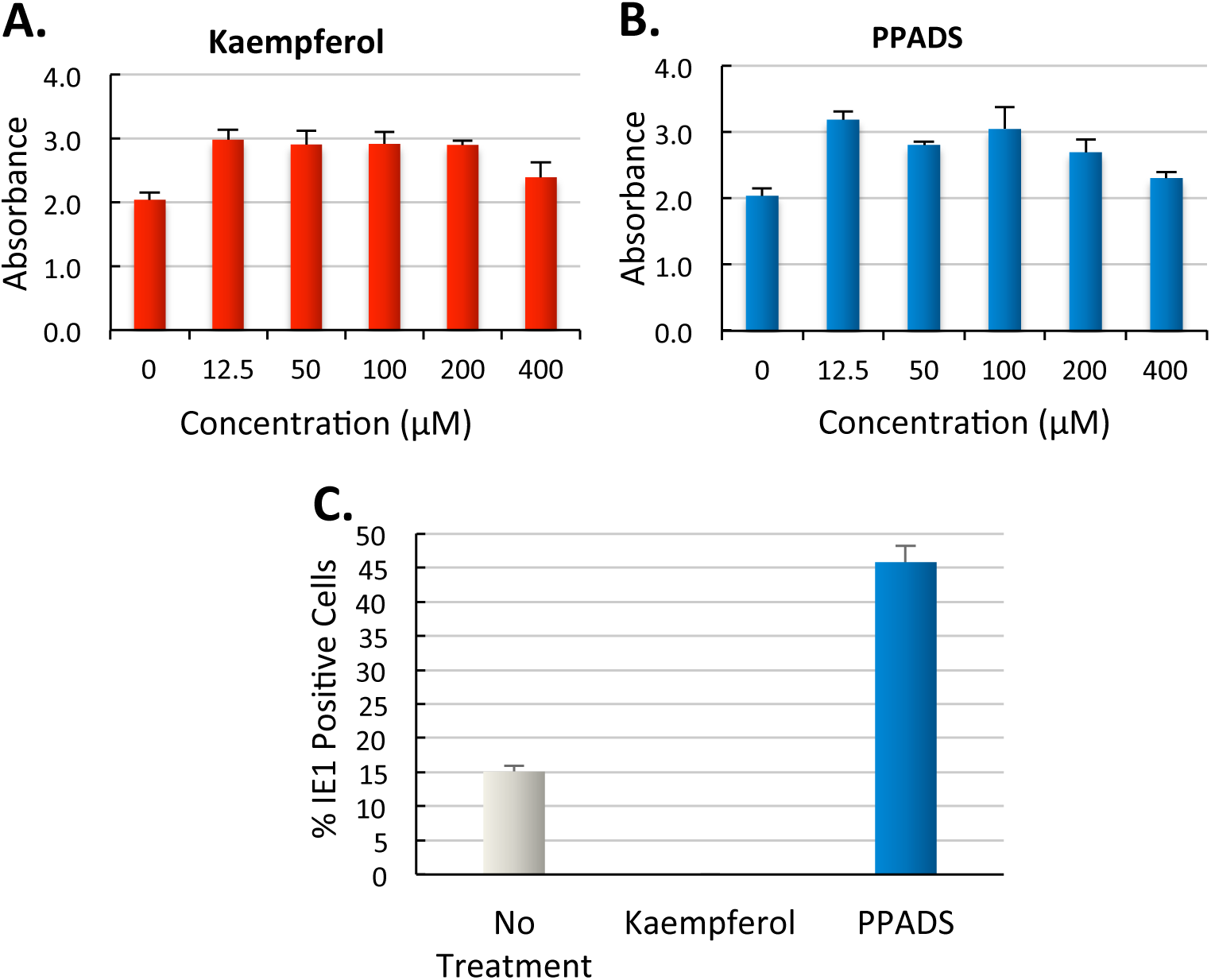
Kaempferol and PPADS have opposite effects on HCMV yield. (A, B) Drug toxicity assays. HFFs were treated with various concentrations of (A) kaempferol and (B) PPADS. The media with drugs was replenished every 24 h. At 96 h, cell viability was measured using the CellTiter 96^®^ AQ_ueous_ Assay. Results are shown as average absorbance from two biological replicates and error bars represent SEM. **(C) Drugs modulate the yield of HCMV**. HFFs were infected with TB40/E-GFP virus (MOI=3) or mock infected for 2 h and treated with either kaempferol (50 μM) or PPADS (50 μM). The drugs were replaced every 24 h until media samples were collected at 96 hpi. Viral yield was determined by monitoring expression of IE1 protein at 24 h after infection. Results were averaged between two biological replicates and error bars represent SEM.

### Inhibiting P2Y2 and P2X5 purinergic receptors does not affect the efficiency of viral entry into fibroblasts

We next aimed to elucidate the point during infection at which the two purinergic receptors play a role. A possible explanation for the observed changes in viral yield upon inhibiting the activity or expression of P2Y2 and P2X5 is that the cell surface purinergic receptors may function in the initial entry of the virus into fibroblasts. Although this notion did not fit well with the kinetics of receptor expression following HCMV infection (Fig. 2), purinergic signaling was shown to be involved in the entry step for HIV-1 infection [17, 59]. Therefore, we tested this possibility using both pharmacological antagonists and siRNA-mediated expression knockdown. First, we assessed the efficiency of viral entry into cells that have been pre-treated with kaempferol or PPADS compared to solvent controls. Cells were treated with drugs and after 1 h, cells were washed and HCMV-infected at a multiplicity of 1 TCID_50_/cell for 1 h at 4°C to allow virus attachment to the cell surface. Then, the viral particles that did not attach were removed and cells were incubated at 37°C, allowing the attached virus to penetrate the cells. After 24 h, the cells were fixed and immunostained for IE1 protein. Pre-treatment with either drug did not significantly alter the percentage of cells expressing IE1 protein compared to solvent controls (Fig. 6A). To confirm these results, we assessed the efficiency of viral entry in cells transfected with siP2Y2, siP2X5 or siSc as a control. Cells were transfected with the appropriate siRNA for 24 h before HCMV infection at a multiplicity of 1 TCID_50_/cell. After 1 h at 4°C, unattached virions were removed and cultures were shifted to 37°C. One hour later, samples were collected and viral DNA copy number was quantified by qPCR. Again, results showed no significant differences in fold change of viral DNA copies between the siP2Y2 or siP2X5 transfected cells and the siSc controls (Fig. 6B). We conclude that inhibiting P2Y2 and P2X5 activity or expression does not affect viral entry into HFFs.

**FIG 6.**
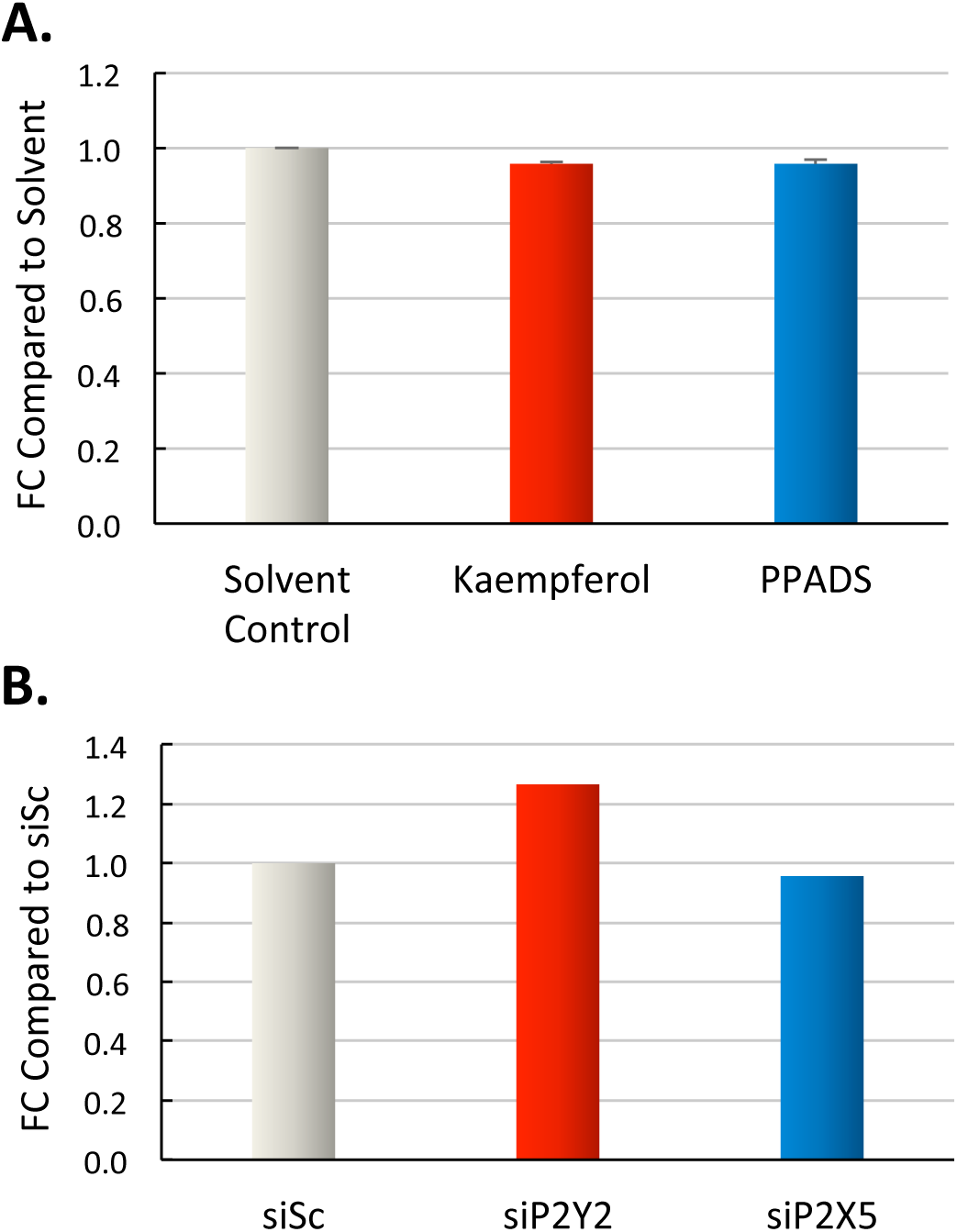
P2Y2 and P2X5 receptors do not affect HCMV entry into HFFs. HFFs were either (A) pre-treated with either kaempferol (50 μM) or PPADS (50 μM), or solvent control for 1 h, or (B) transfected with siP2Y2, siP2X5, or siSc as a control for 24 h. Then, they were infected with TB40/E-GFP virus (MOI=1) or mock-infected for 1 h at 4°C. Cells were washed with citrate buffer and incubated at 37°C. Viral entry was assayed by either (A) immunostaining for IE1 protein at 24 hpi or (B) qPCR quantification of intracellular viral DNA at 1 hpi. Results are reported as fold change (FC) compared to the solvent control or siSc control conditions. Data for two technical replicates were averaged and error bars represent SEM.

### Inhibiting P2Y2 reduces the accumulation of viral transcripts

To probe when and how the cellular P2Y2 receptor might affect viral gene expression subsequent to virus entry, we analyzed viral gene expression in siSc-and siP2Y2-treated cells at various times during the viral replication cycle. qRT-PCR was used to quantify transcript levels for representatives of the three main classes of viral genes. The data show that infection in cells transfected with siP2Y2 resulted in a reduction of immediate early (UL123, UL122), early (UL26, UL54), and late (UL69, UL82, UL99) viral transcripts compared to siSc controls (Fig. 7A-B, D-H). Viral RNAs were most dependent on P2Y2 during the late phase of infection, consistent with the timing of P2Y2 RNA accumulation (Fig. 2). Intriguingly, one viral transcript, immediate early UL37x1, was expressed at higher levels in cells transfected with siP2Y2 than in control cells (Fig. 7C). UL37x1 has been implicated in the mobilization of intracellular Ca^2+^ during infection [28, 30].

**FIG 7.**
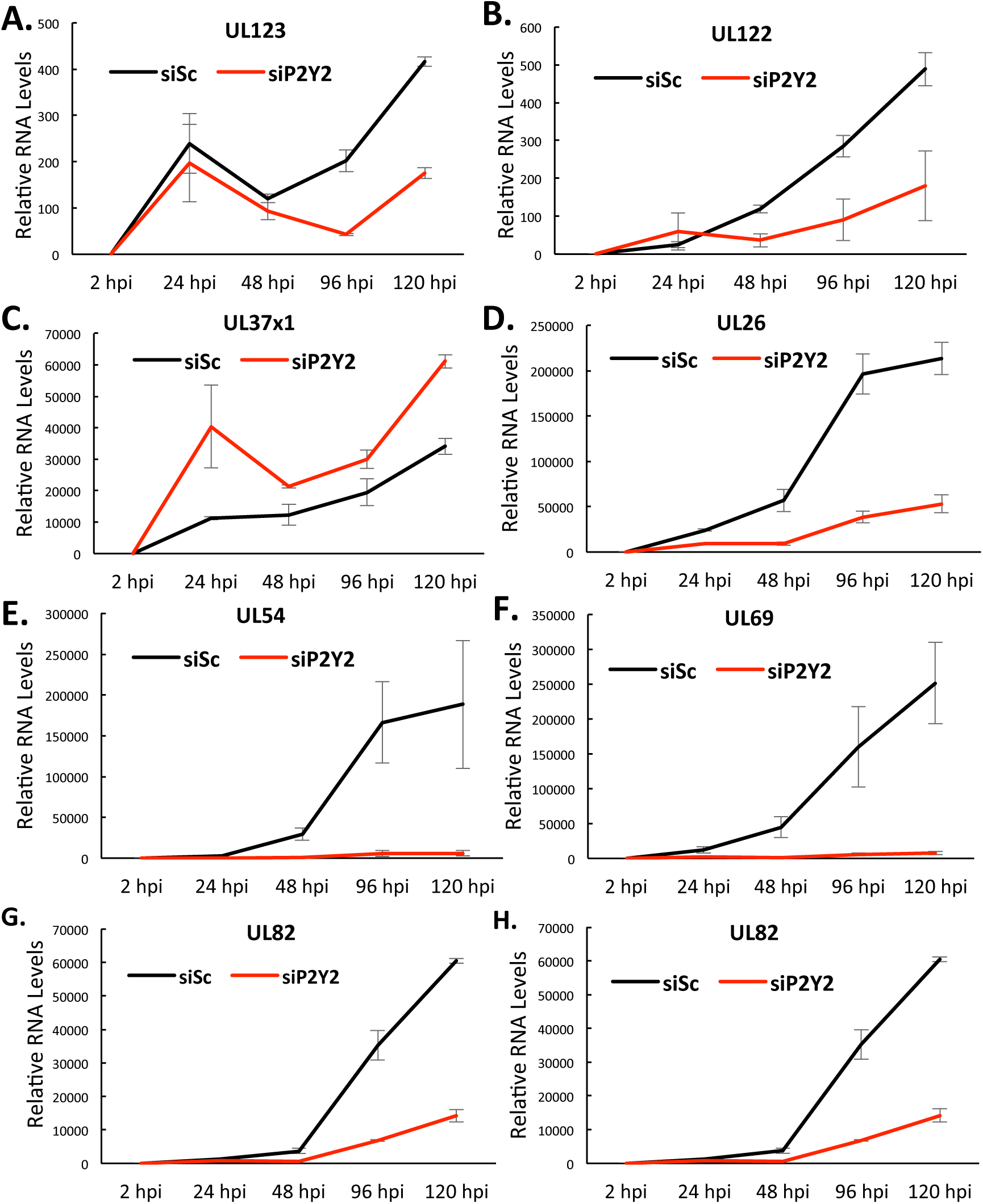
P2Y2 receptor regulates levels of viral gene expression. HFFs were transfected with siP2Y2 or siSc as a control. After 24 h, cells were infected with TB40/E-GFP virus (MOI=3) or mock infected, and RNA samples were collected after various time intervals. Viral transcript levels were determined by qRT-PCR with GAPDH serving as an internal control. The data are shown as fold change compared to transcript levels at 2 hpi, calculated from two technical replicates. Error bars represent SEM.

### P2Y2 receptor affects the efficiency of viral DNA synthesis

One of the strongest inhibitory effects of P2Y2 knockdown was seen on the accumulation of RNA coding for the viral DNA polymerase, UL54 (Fig. 7E). Given its direct role in viral DNA replication [60], we tested whether P2Y2 could have an impact on viral DNA (vDNA) accumulation. Total DNA was isolated from HCMV-infected control and P2Y2-deficient cells and vDNA copy numbers were measured by qPCR using primers specific to UL123. There were about 5-fold fewer copies of vDNA in P2Y2-deficient cells than in control cells (Fig. 8A). This supports the hypothesis that P2Y2 inhibition prevents viral DNA accumulation by blocking viral DNA synthesis. To confirm this result, we also compared the vDNA copy numbers in released viral particles between siP2Y2-treated cells and siSc-treated controls. Similarly, when viral genomic DNA was isolated from viral particles and quantified, there were also nearly 5 times fewer copies of vDNA released from P2Y2-deficient cells compared to control cells (Fig. 8B). To confirm that lower viral DNA synthesis is the predominant explanation for the decrease in infectious particles found in the media collected from P2Y2-deficient cells (Fig. 4B, C and 5C), we decided to test the alternative hypothesis that progeny released from P2Y2 knockdown cells are less infectious by investigating if there are any effects of P2Y2 inhibition on the infectivity of virions. Accordingly, the ratios of vDNA copy number to infectious unit were calculated, and there was no significant difference between the siP2Y2-treated cells and siSc controls (Fig. 8C). Therefore, the reduction in HCMV yield observed in the P2Y2 knockdown condition is caused by reduced viral DNA synthesis and release of viral particles, and not by reduced infectivity of the virions released from P2Y2-deficient cells. Taken together, it appears that P2Y2 is critical for viral DNA synthesis. Inhibiting the P2Y2 receptor reduces viral DNA synthesis and causes a drastic reduction in viral titer.

**FIG 8.**
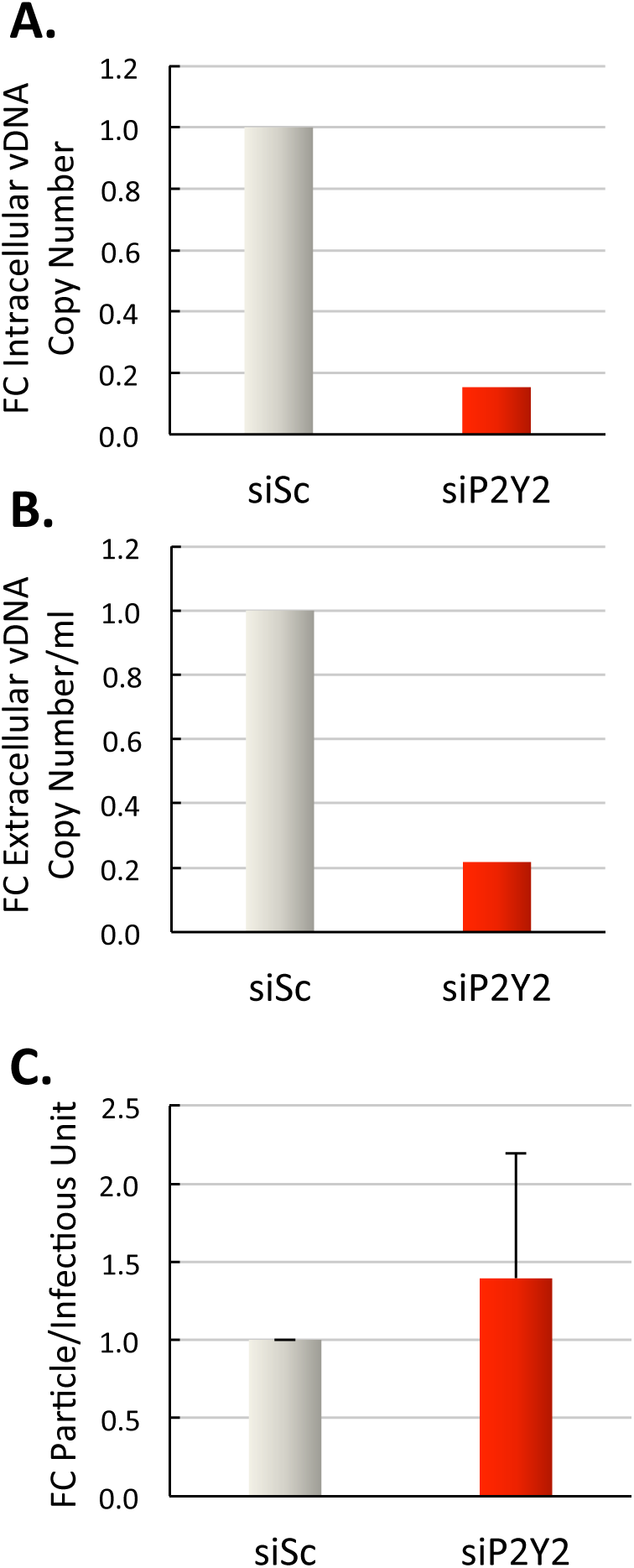
P2Y2 receptor affects viral DNA accumulation. HFFs were transfected with siP2Y2 or siSc as a control. After 24 h, cells were infected with TB40/E-GFP virus (MOI=3) or mock infected. (A) **P2Y2 KD reduces intracellular vDNA**. At 96 hpi, total DNA was isolated from siSc-or siP2Y2-treated, HCMV-infected cells and the level of intracellular viral DNA was measured by qPCR. **(B) P2Y2 KD reduces extracellular vDNA**. At 96 hpi, total DNA was isolated from media of siSc-or siP2Y2-treated, HCMV-infected cells and the level of extracellular viral DNA was measured by qPCR. **(C) P2Y2 KD does not change the particle-to-infectious unit ratio**. The infectivity of virus in media collected at 96 hpi was titered and viral DNA was isolated from virions present in the media and quantified by qPCR to calculate a viral DNA copy number-to-infectious unit ratio. The results are depicted as fold change of this ratio. Data were averaged between two biological replicates and error bars represent SEM.

### P2Y2 and UL37x1 cooperate to regulate intracellular Ca^2+^ during infection

One of the most striking findings in our analysis of the role of purinergic receptors on viral gene expression during HCMV infection is the fact that of all the viral genes studied, only one was upregulated in the siP2Y2-treated cells compared to siSc-treated control cells (Fig. 7C). This was the transcript of the viral immediate early UL37x1 gene. Previously, UL37x1 has been shown to be involved in the release of Ca^2+^ from SER stores to increase cytosolic Ca^2+^ concentrations during infection [28, 30]. Interestingly, P2Y2 is a G-protein coupled receptor that, when activated, leads to the downstream hydrolysis of PIP_2_ into IP_3_, which is also known to cause the release of Ca^2+^ ions from SER stores [11, 61]. Because of the potential involvement of P2Y2 in Ca^2+^ homeostasis following infection, we investigated the effects of inhibiting P2Y2 expression on intracellular Ca^2+^ levels. To do so, fibroblasts were infected with either wild-type AD*wt* virus or AD*sub*UL37x1 virus lacking the UL37x1 gene at a multiplicity of 1 TCID_50_/cell or mock infected, and cultures were treated 2 h later with kaempferol or solvent control. Using a multiplicity of 1 TCID_50_/cell allowed us also to measure Ca^2+^ levels in infected and uninfected cells in the same experimental well. At 20 hpi, we used the fluorescence-based fluo-4 AM calcium assay to compare the changes in intracellular Ca^2+^ concentrations. We determined that wild type virus caused ~2.5-fold increase in Ca^2+^ levels compared to uninfected cells (Fig. 9A). We postulated that if P2Y2 contributes to elevated levels of Ca^2+^ in infected cells, then we should see a partial decrease of Ca^2+^ levels in cells either infected with AD*wt* and treated with kaempferol or infected with AD*sub*UL37x1. Instead, we saw that in both instances, Ca^2+^ was reduced almost to the level found in uninfected cells (Fig. 9A). Interestingly, Ca^2+^ levels did not decrease further in cells infected with AD*sub*UL37x1 and treated with kaempferol. Therefore, these data imply that P2Y2 and pUL37x1 work together to elevate Ca^2+^ levels during HCMV infection and that the lack of either component prevents cells from increasing intracellular calcium levels during infection.

**FIG 9.**
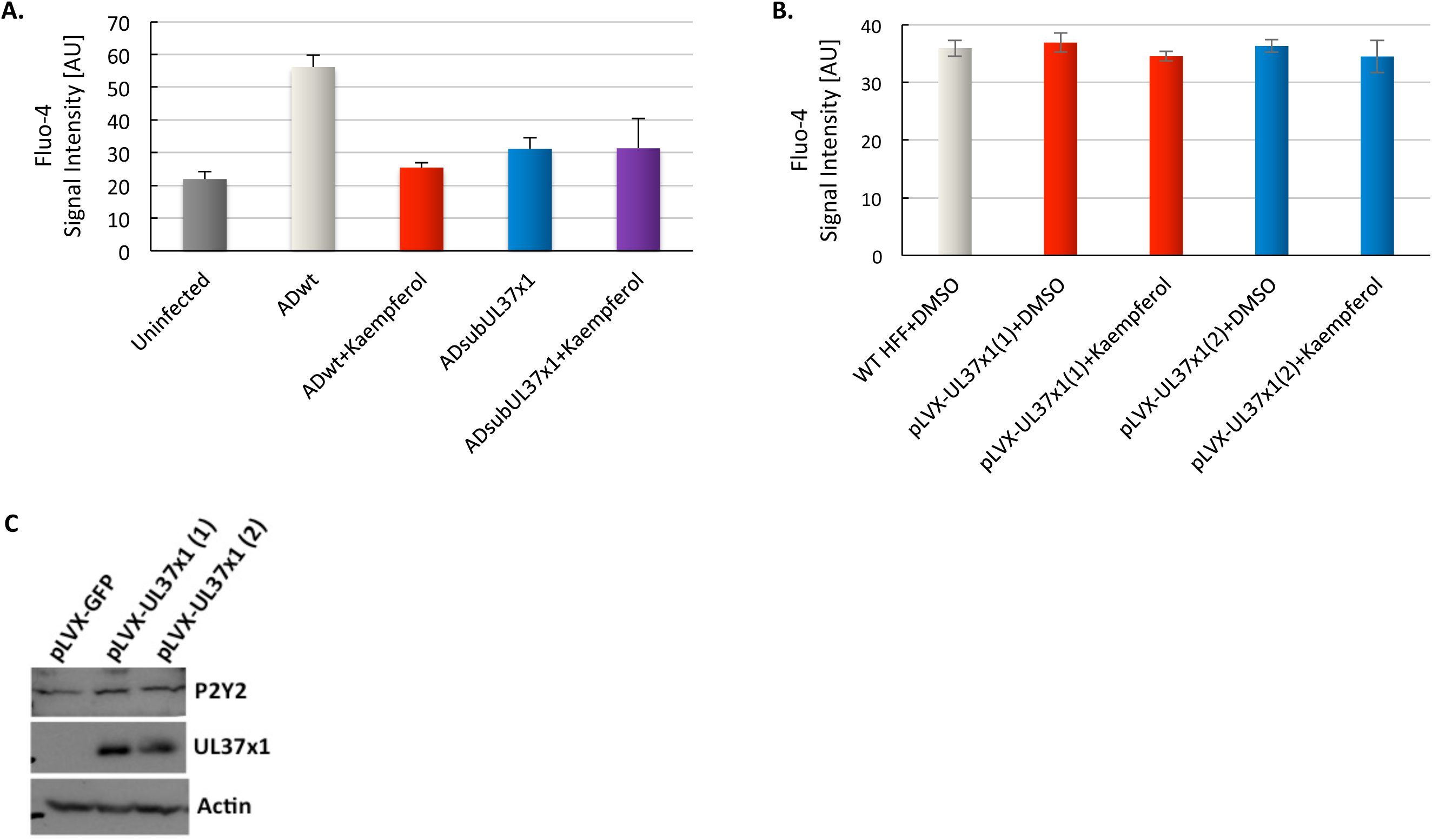
P2Y2 and UL37xl work cooperatively to regulate Ca^2+^ in HCMV-infected cells. **(A) Effects of kaempferol on Ca^2+^ levels in AD*wt* vs. AD*sub*UL37x1 infected cells**. HFFs were infected with either AD*wt* or AD*sub*UL37x1 (MOI=l) or mock infected for 2 h and treated with kaempferol (50 μM) or solvent control. Fluo-4 AM dye was applied at 20 hpi. **(B) Effects of kaempferol on Ca^2+^ levels in wild type vs. UL37xl overexpressing cells**. Wild type HFF (WT HFF) cells or HFF cells overexpressing UL37xl [pLVX-UL37x1(1) and pLVX-UL37xl(2)] were treated with kaempferol (50 μM) or solvent control. Fluo-4 AM dye was applied 1 h later. Pictures of cells were taken using the Operetta High-Content Imaging System and the fluorescent signal was measured using ImageJ software based on 10 cells per experimental arm. Results are presented as the intensity of fluorescent signal in arbitrary units [AU]. **(C) P2Y2 and UL37xl protein in UL37xl overexpressing cells**. Protein samples were collected from two clones of MRC5 fibroblasts overexpressing UL37xl [pLVX-UL37xl (1) and pLVX-UL37xl (2)] and control cells expressing GFP Proteins were separated on SDS-PAGE and Western blot analysis was performed using antibodies recognizing P2Y2, UL37xl and actin as a loading control.

To further test this hypothesis, we investigated the effect of UL37x1 expression on Ca^2+^ homeostasis in HFF cells without the background of infection. For this purpose, a UL37x1-expressing lentiviral pLVX-EF1 plasmid was created. HFF cells were transduced with the vector, and subjected to puromycin selection. Two cellular clones were obtained [pLVX-UL37x1(1) and pLVX-UL37x1(2)] that exhibited stable expression of pUL37x1. The cells were treated with kaempferol or solvent control for 1 h. Then, Ca^2+^ levels were measured using the fluo-4 AM assay and compared to levels in wild type HFF cells. No significant changes in Ca^2+^ levels were observed between samples (Fig. 9B), which supports our conclusion that increased expression of pUL37x1 in the absence of P2Y2 overexpression is not sufficient to stimulate the robust increase in calcium seen in HCMV-infected cells.

As a control, cells transduced with pLVX-UL37x1 or pLVX-GFP plasmids were analyzed by Western blot assay for the level of pUL37x1 expression. The assay confirmed the pUL37x1 overexpression only in pLVX-UL37x1 plasmid-transduced HFF cells (Fig. 9C). Because inhibiting the P2Y2 receptor activity decreases the extent to which intracellular Ca^2+^ is mobilized during infection, we propose that P2Y2 signaling plays a critical role in controlling intracellular Ca^2+^ during HCMV infection.

Since previous studies as well as our current study have shown that the viral pUL37x1 protein is involved in the mobilization of intracellular Ca^2+^ from calcium stores in the SER and knowing that P2Y2 expression was upregulated early in infection, we aimed to further elucidate the relationship between the viral UL37x1 and cellular P2Y2. Specifically, we assessed whether the upregulation of P2Y2 expression during viral infection may be a downstream effect of the immediate early kinetics of UL37x1 expression [62]. Since the upregulation of P2Y2 transcript is most evident around 48hpi, it is conceivable that the viral pUL37x1 protein increases the expression of P2Y2 and that increased P2Y2 activity further increases intracellular Ca^2+^ levels for the remainder of the infection. In order to test this idea, HFFs were infected with either the AD*sub*UL37x1 virus or the wild type AD*wt* virus. Samples were collected at 24 hpi. P2Y2, UL54, UL123 and UL37x1 transcript levels were quantified by qRT-PCR. If the hypothesis that P2Y2 upregulation is dependent on pUL37x1 expression were true, then P2Y2 transcripts would be significantly lower in cells infected with AD*sub*UL37x1 than AD*wt* virus. However, although P2Y2 expression was appeared somewhat lower in cells infected with AD*sub*UL37x1 compared to those infected with AD*wt*, the difference was not significant (Fig. 10A). Interestingly, it was also noted that unlike siP2Y2-treated cells, AD*sub*UL37x1-infected cells did not exhibit lower UL54 RNA levels compared to AD*wt*-infected cells (Fig. 10B). The same expression levels of UL123 in cells infected with either AD*wt* or AD*sub*UL37x1 confirm that the cells received an equal amount of virus (Fig. 10C). As expected, UL37x1 expression was only evident in AD*wt*-and not in AD*sub*UL37x1-infected cells (Fig. 10D). Both clonal populations of cells expressing pUL37x1 were characterized by a robust expression of pUL37x1, but there was only a minimally higher level of P2Y2 protein in these cells compared to controls expressing GFP (Fig. 9C). Therefore, these data suggest that viral pUL37x1 is likely not a significant upstream activator of P2Y2 expression.

**FIG 10.**
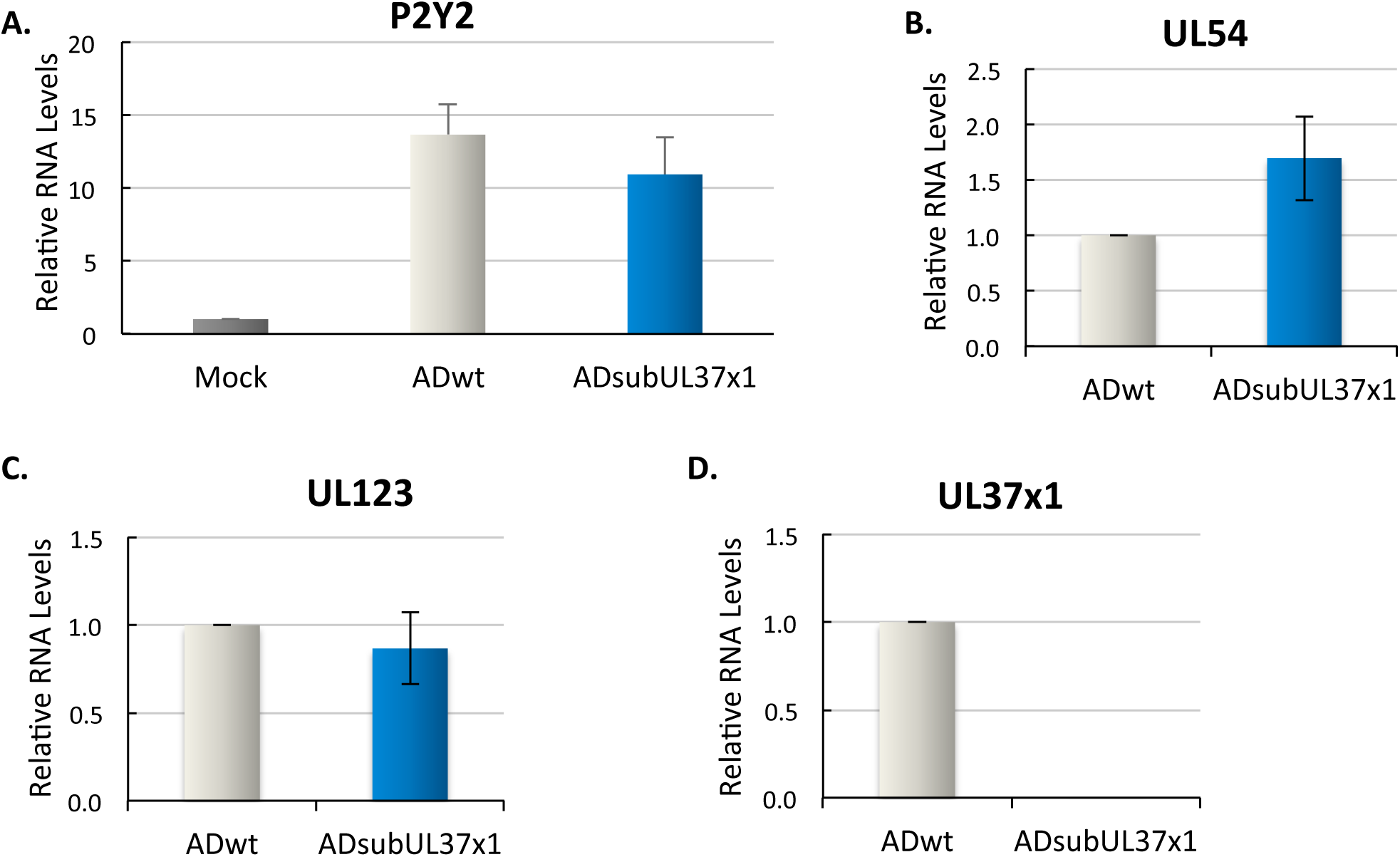
P2Y2 overexpression in HCMV-infected HFFs is not regulated by viral UL37xl. sHFFs were infected with either AD*wt* or AD*sub*UL37x1 virus (MOI=3) or mock-infected for 2 h. Samples were collected at 24 hpi. (**A**) P2Y2, (**B**) UL54, (**C**) UL123 and (**D**) UL37x1 transcript levels were determined by qRT-PCR with GAPDH serving as an internal control. Results are shown as fold change compared to mock-infected or AdwMnfected controls. Data are averaged across three biological replicates and error bars represent SEM.

Taken together, these findings suggest that both cellular P2Y2 and viral pUL37x1 cooperate to modulate intracellular Ca^2+^ within HCMV-infected cells.

## Discussion

The purpose of this study was to investigate the role of purinergic signaling during HCMV infection in human fibroblasts. RNA-Seq analysis identified several members of the purinergic receptor network whose expression was modulated upon infection (Table 2). qRT-PCR analysis confirmed that infection increased the cellular P2Y2, P2X5, ENPP4 and ENTPD2 transcripts. Elevated expression of purinergic receptors during HCMV infection has been reported previously. In myeloid cells, HCMV infection increases the expression of the P2Y5 receptor [63]. In endothelial cells, HCMV infection has been associated with higher levels of the P2Y2, P2Y1, and P2X7 receptors, as well as ecto-5’nucleotidase CD73 [25, 26]. Yet, the impact of these altered expression patterns on the efficiency of HCMV infection has not been investigated. We now report that siRNA targeting ENPP4 and ENTPD2 transcripts did not affect viral yield (Fig, 4B). In contrast, inhibiting the activity or expression of P2Y2 and P2X5 receptors had a robust effect on the amount of infectious progeny released from infected cells (Fig. 4B, C). These findings suggest that, at least in fibroblasts, purinergic signaling through the P2Y2 and P2X5 receptors is important in HCMV infection independent from the activity of ectonucleotidases.

Inhibiting P2Y2 and P2X5 by using pharmacological compounds or siRNA technology drastically altered HCMV yield. It is noteworthy that, even though both receptors are activated by ATP, they had completely opposite effects on the production of HCMV progeny. Compared to control treated cells, infection in P2Y2-deficient cells led to a drastic reduction (~95% lower) in the number of infectious virions released (Fig. 4B, C), which demonstrated that P2Y2 supports HCMV replication. On the other hand, infection in P2X5-deficient cells resulted in enhanced (4x higher) viral release compared to control treated cells (Fig. 4B, C), suggesting that P2X5 antagonizes HCMV replication.

Our analysis of P2Y2 and P2X5 expression kinetics supported the view that these receptors are independently regulated and have different functions during HCMV infection. While active viral gene expression is necessary for the upregulation of P2Y2 expression (Fig. 3A), it is not for the increased expression of P2X5 (Fig. 3B). Moreover, P2Y2 gradually increased in expression from 2 to 96 hpi, as seen at both RNA and protein levels (Fig. 2A, C), P2X5 expression increased up to 48 hpi (RNA) or 72 hpi (protein) and then decreased immediately afterwards (Fig. 2B, C). The gradual increase in P2Y2 expression is consistent with a supportive role for the receptor throughout the viral replication cycle. Conversely, the initial spike in the levels of the inhibitory P2X5 receptor might be antagonized by a virus-mediated mechanism that suppresses its expression later after infection.

How do the P2Y2 and P2X5 receptors modulate HCMV replication? The P2X5 purinergic receptor is a non-selective cation channel that is gated by extracellular ATP. It is typically found on cells of the skin, gut, bladder, thymus, skeletal muscle, and spinal cord and is thought to play important roles in cell differentiation [64]. For instance, it has been shown to be involved in the differentiation of skin keratinocytes and mucosal epithelial cells during turnover in the gut and bladder [19, 65]. P2X5 receptors have also been observed to play various roles in regulating osteoblastic differentiation and proliferation, triggering differentiation of skeletal muscle satellite cells, and the differentiation of human fetal epidermis [66, 67]. Although the expression of P2X5 receptors have been associated with differentiating and proliferating cell types, the exact mechanism by which its signaling affects these cellular processes is still unclear. Thus it is difficult to speculate on how a block to its activity might influence HCMV replication.

P2Y2 is a G-protein coupled receptor that can be activated by extracellular ATP and UTP. HCMV infection of P2Y2 knockdown cells exhibited reduced expression of most viral genes tested, including UL54, the viral DNA polymerase catalytic subunit (Fig. 7E). Reduced pUL54 levels would be expected to reduce key downstream events in the viral replication cycle. Indeed, P2Y2-deficient cells exhibited reduced accumulation of viral DNA (Fig. 8A) and late viral RNAs (Fig. 7F-H). Although it is likely that the receptor has additional effects on HCMV replication, its effect on viral DNA accumulation, followed by inhibition of late RNAs that depend on active DNA replication for their synthesis, can explain the low viral yield from P2Y2-deficient cells.

There are several plausible mechanisms by which purinergic signaling via the P2Y2 receptor can affect viral DNA replication. It could affect DNA replication indirectly, through inhibition of an upstream viral function that we have not yet measured, or it could act directly at DNA replication. One potential mechanism is through p38 mitogen-activated protein kinases (MAPKs). Extracellular ATP and UTP can activate the MAPK kinase (MKK) 3/6-p38-MAPK cascade via the P2Y2 receptor in glomerular mesangial cells [68]. p38 activation results from increased activity of MKK3/6, which are downstream effectors of P2Y2 signaling [69]. It has been reported that active p38 plays a critical role in HCMV viral DNA replication in infected human fibroblasts [69]. Thus, inhibiting P2Y2 could decrease MKK3/6 activity, which might then reduce p38 activation and viral DNA replication. This role for P2Y2 action within infected cells remains speculative, because the exact mechanism by which p38 regulates viral DNA replication via the UL54 polymerase has not yet been characterized.

In addition to its impact on p38 MAPK, the P2Y2 receptor has a well-established role in regulating cellular calcium levels [15, 16, 23]. Extracellular nucleotides act via the P2Y2 receptor to induce intracellular Ca^2+^ mobilization from SER in a variety of cell types [70–72]. The increase in intracellular Ca^2+^ can be blocked by PLC inhibitors and by low molecular weight heparin, indicating the involvement of IP_3_-sensitive intracellular Ca^2+^ stores, which is known to be downstream of P2Y2-mediated signaling [73]. As calcium homeostasis was found to be critical for an efficient HCMV infection [27, 28], it was intriguing to speculate that P2Y2 inhibition may have important consequences for HCMV replication due to disruption of intracellular calcium regulation. Treating infected cells with drugs that disrupt SER Ca^2+^ homeostasis inhibits the production of infectious progeny due to retarded accumulation of late gene products [27], patterns that were observed during HCMV infection in P2Y2 knockdown cells.

The first indication that P2Y2-dependent Ca^2+^ regulation may be important during HCMV infection came from analyzing differential expressions of viral genes in wild type cells and cells treated with siP2Y2. Among all tested viral transcripts only UL37x1 RNA levels were upregulated in P2Y2-deficient cells (Fig. 7C). It is known that the HCMV immediate early protein pUL37x1 induces the mobilization of Ca^2+^ from the SER to the cytosol [28]. In addition, the Ca^2+^-dependent protein kinase, PKCα, is activated following infection, leading to the production of large (1-5 μm diameter) cytoplasmic vesicles late after infection [30]. The presence of these vesicles correlated with the efficient accumulation of enveloped virions. Regulating intracellular Ca^2+^ levels may also influence viral DNA replication. Ca^2+^ can induce the activity of Ca^2+^/calmodulin-dependent protein kinase kinase (CaMKK), which activates AMPK. AMPK activity has been shown to be necessary for HCMV DNA synthesis and the expression of viral late genes [33, 34]. In addition, it has been found that extracellular nucleotides stimulate the PI3-K/Akt pathway through P2Y2-mediated signaling involving Ca^2+^ influx, CaM, and Src [74]. It was previously reported that treating infected fibroblasts with an inhibitor of PI3-K activity caused inhibition of viral DNA replication and a 4-log decrease in viral titers [36]. Although intriguing, it is more likely that P2Y2-mediated inhibition of viral UL54 is independent from its effects on intracellular Ca^2+^. Cells infected with AD*sub*UL37x1, which lacks the UL37x1 gene, show reduced intracellular Ca^2+^ levels (Fig. 9A), but, nevertheless, do not show reduced UL54 RNA levels (Fig. 10B). Hence the effect P2Y2 on viral DNA accumulation may be distinct from its effects on intracellular Ca^2+^ concentrations.

To broaden our understanding on a potential role of P2Y2 in Ca^2+^ levels and the relationship between P2Y2 and UL37x1 during infection, we tested the effect of kaempferol, a P2Y2 antagonist, on intracellular Ca^2+^ in cells infected with wild-type HCMV (AD*wt*) versus the pUL37x1-deficient mutant virus (AD*sub*UL37x1). As shown previously [28], AD*wt*-infected cells exhibited increased free Ca^2+^ compared to uninfected cells, whereas AD*sub*UL37x1-infected cells had Ca^2+^ levels similar to those in uninfected cells (Fig. 9A). Interestingly, kaempferol treatment also lowered Ca^2+^ levels in AD*wt*-infected cells to its level in uninfected cells and similar Ca^2+^ levels were measured in kaempferol-treated, AD*sub*UL37x1-infected cells (Fig. 9A). Additionally, when we tested Ca^2+^ levels in uninfected cells that overexpress UL37x1, no elevated levels of Ca^2+^ were observed (Fig. 9B). While P2Y2-deficient cells were characterized by increased expression of UL37x1 (Fig. 7C), neither AD*sub*UL37x1-infected cells nor cells overexpressing UL37x1 affected P2Y2 RNA or protein levels (Fig.10A, 9C). Taken together, these results strongly suggest that P2Y2 has a critical role in regulating Ca^2+^ levels following HCMV infection. Both P2Y2 and pUL37x1 are indispensable for maintaining favorable intracellular Ca^2+^ levels during infection. It is not yet clear how the viral and cellular gene products cooperate in this process.

Aside from disrupting viral DNA replication and Ca^2+^ homeostasis, a third mechanism by which P2Y2 activation may play a role in HCMV infection is through affecting the nuclear egress of the viral capsid. It has been reported that PKC, a downstream effector of P2Y2 signaling, plays an important role in destabilizing the nuclear lamina by phosphorylating several types of nuclear lamin [75]. The phosphorylation and reorganization of the nuclear lamins is an essential part of viral nuclear egress [75]. It is conceivable, then, that a block to nuclear egress of HCMV capsids could contribute to the reduction in infectious virus produced following inhibition of P2Y2 (Fig. 4, 5).

It has been reported that during hypoxic conditions, fibroblasts release ATP, which activates P2Y2 to regulate cellular DNA synthesis via extracellular signal-regulated kinases 1 and 2 (ERK_1/2_) induced Egr-1 activation [14]. Additionally, it has been found that in human cardiac fibroblasts, ATP-mediated activation of the P2Y2 receptor leads to cell cycle progression and cell proliferation [76]. Moreover, P2Y2 activation results in the activation of PKC, which acts in concert with ERK and PI3-K/PKC pathways to induce c-Fos protein expression and HeLa cell proliferation [77]. Since P2Y2 is dramatically overexpressed in HCMV-infected cells, it would be interesting to understand how the expected upregulation of cellular DNA synthesis and cell cycle progression is avoided in HCMV-infected cells in order to maintain the cell cycle arrest characteristic of infection [78, 79].

Signaling via the P2Y2 receptor also may influence cell migration. P2Y2 activation increases MCF-7 breast cancer cell migration via the MEK-ERK_1/2_ signaling pathway [80]. Specifically, extracellular ATP can activate MAPKs through the P2Y2/PLC/PKC/ERK signaling pathway to induce the translocation of ERK_1/2_ into the nucleus [13]. Also, fibroblasts appear to require PKC activation, which is downstream of P2Y2 signaling, in order to respond to hyaluronan stimulation with increased locomotion [81]. Taken together, it is conceivable that the overexpression of P2Y2 receptors during HCMV infection may lead to increased migration of the host cell. This not only has potential to facilitate dissemination of the virus within its infected host, it may also contribute to the likely oncomodulatory activity of HCMV [82].

In summary, our results show that purinergic signaling through the P2Y2 and P2X5 receptors plays a critical role within infected cells and set the stage for additional investigations on the impact of this receptor family in HCMV biology. Further, our observation that a pharmacological block to P2Y2 dramatically reduces viral yield raises the possibility that P2Y2 antagonists, if well tolerated, could prove to be attractive candidates for new HCMV therapies.

## Acknowledgements

This work was supported by a grant from the National Institutes of Health (AI112951). M.N. was supported by a fellowships from the American Cancer Society (PF-14-116-01-MPC).

